# Anti-Insulin receptor antibodies improve hyperglycaemia in a mouse model of human insulin receptoropathy

**DOI:** 10.1101/2020.03.20.999771

**Authors:** Gemma V Brierley, Hannah Webber, Eerika Rasijeff, Sarah Grocott, Kenneth Siddle, Robert K Semple

**Affiliations:** The University of Cambridge Metabolic Research Laboratories, Wellcome Trust-MRC Institute of Metabolic Science, Addenbrooke’s Hospital, Cambridge, CB2 0QQ, UK; MRC Disease Model Core, Metabolic Research Laboratories, Wellcome Trust-MRC Institute of Metabolic Science, Addenbrooke’s Hospital, Cambridge, CB2 0QQ, UK; University of Edinburgh Centre for Cardiovascular Science, Queen’s Medical Research Institute, Little France Crescent, Edinburgh, EH16 4TJ

## Abstract

Loss-of-function mutations in both alleles of the human insulin receptor gene (INSR) cause extreme insulin resistance (IR) and usually death in childhood, with few therapeutic options. Bivalent anti-receptor antibodies can elicit insulin-like signaling by mutant INSR in cultured cells, but whether this translates into meaningful metabolic benefits *in vivo*, where dynamics of insulin signaling and receptor recycling are more complex, is unknown. To address this we adopted a strategy to model human insulin receptoropathy in mice, using *Cre* recombinase delivered by adeno-associated virus to knock out endogenous hepatic *Insr* acutely in floxed *Insr* mice (L- IRKO+GFP), before adenovirus-mediated ‘add-back’ of wild-type (WT) or mutant human *INSR*. Two murine anti-INSR monoclonal antibodies, previously shown to be surrogate agonists for mutant INSR, were then tested by intraperitoneal injections. As expected, L-IRKO+GFP mice showed glucose intolerance and severe hyperinsulinemia, and this was fully corrected by add-back of WT but neither D734A nor S350L mutant INSR. Antibody injection improved glucose tolerance in D734A INSR-expressing mice and reduced hyperinsulinemia in both S350L and D734A INSR-expressing animals, and did not cause hypoglycemia in WT INSR-expressing mice. Antibody treatment also downregulated both wild-type and mutant INSR protein, attenuating its beneficial metabolic effects. Anti-INSR antibodies thus improve IR in an acute model of insulin receptoropathy, but these findings imply a narrow therapeutic window determined by competing effects of antibodies to stimulate receptors and induce their downregulation.

**One Sentence Summary:** Bivalent anti-insulin receptor antibodies improve glycaemic control, but downregulate receptor expression, in a novel mouse model of lethal human insulin receptoropathy.

## Introduction

Insulin exerts metabolic and growth promoting effects that are essential for life via a homodimeric plasma membrane receptor tyrosine kinase. Insulin binding to extracellular sites induces alterations in receptor structure that promote *trans* autophosphorylation of tyrosine residues on intracellular beta subunits. This in turn leads to recruitment and phosphorylation of insulin receptor substrate (IRS) proteins, and thence activation of a signalling network, critically including PI3K/AKT and RAS/MAPK pathways*(1)*.

Attenuated glucose-lowering by insulin *in vivo* is referred to as insulin resistance (IR), and is a core feature of the metabolic syndrome in humans. IR is closely associated with obesity, type 2 diabetes, an abnormal blood lipid profile that promotes atherosclerosis, fatty liver and reduced fertility, but its molecular and cellular basis is not fully elucidated*(2)*. Several severe forms of IR are known where the precise cause is established, however. The most extreme of these are caused by bi-allelic insulin receptor *(INSR*) mutations, and are clinically described as Donohue or Rabson–Mendenhall syndromes (OMIM#246200 or #262190). Death is usual within the first 3 years in Donohue syndrome, while in Rabson Mendenhall syndrome mortality in the second or third decades is common. There is a major unmet need for novel approaches to circumvent impaired function of mutant receptors. Some INSR mutations impair receptor processing and thus cell surface expression, however many mutant INSR are well expressed at the cell surface, but exhibit impaired insulin binding and/or impaired signal transduction*(3)*. This affords the opportunity to activate the mutant receptors using surrogate ligands, and observations of the genetic spectrum of receptoropathy suggest that even modest activation is likely to elicit meaningful metabolic effects*(2)*.

It was first demonstrated in the 1980s that crosslinking of insulin receptor homodimers by bivalent antibodies could elicit signalling responses*(4)*, and in the early 1990s the principle that insulin receptors harbouring disease-causing mutations could also be partly activated by antibodies was provided for two mutations, one in a cell culture model and the other in solubilised form*(5, 6)*. With therapeutic humanised monoclonal antibodies now well established as treatments both for cancer and non-cancer indications*(7)*, interest in biological therapies targeting the INSR has recently rekindled. Inhibitory INSR antibodies are now in Phase 1 human trials*(8)* while stimulatory antibodies have been shown to ameliorate diabetes in rodents*(9–11)* and primates*(12)*. Given the high clinical need in recessive insulin receptoropathy, we previously assessed the effect of monoclonal anti-INSR antibodies*(13–17)* on a series of disease-causing mutant INSRs in cell culture models, corroborating and extending prior findings by demonstrating an action of antibodies against a panel of mutant receptors*(18)*.

Whether the stimulation of mutant receptors by anti-receptor antibodies that is observed biochemically after acute exposure of cells in culture will be sustained and metabolically beneficial *in vivo* has not yet been addressed. A specific concern relates to the documented effect of naturally occurring anti-INSR autoantibodies, which are partial agonists when tested acutely on cellular models, to downregulate INSR signalling when present at high concentrations *in vivo*, inducing acquired severe IR, known as “type B” IR*(19)*. Such an effect has not been assessed in preclinical testing of anti-INSR antibodies reported to date*(9–12)*, but is a critical concern in efforts to develop safe, efficacious biological therapies targeting the INSR.

To address these issues we have now generated a novel model of human insulin receptoropathy restricted to mouse liver, based on adenoviral overexpression of human wild-type or mutant INSR in the liver after *cre*- mediated knockout of endogenous murine *Insr*. Using this model we assessed the effect of two anti-human INSR monoclonal antibodies previous tested in cell culture on metabolic endpoints and receptor expression.

## Results

### Creation of a flexible murine model of human insulin receptoropathy

We first set out to generate a “humanized” mouse model of insulin receptoropathy (Figure 1A). This was necessary as the previously described anti-human INSR monoclonal antibodies to be tested (83-7 and 83-14, which bind distinct epitopes on the receptor α-subunit) were raised in mice and do not bind rodent receptor*(14)*. Eight-week old *Insr*^*loxP/loxP*^ mice were injected *via* the tail vein with an adeno-associated virus (AAV) containing *Cre* recombinase under control of a liver specific promoter (thyroid hormone-binding globulin). This deleted endogenous hepatic *Insr*, generating liver insulin receptor knock out (L-IRKO) mice. As described previously*(20)* this approach avoids the compensatory, secondary responses seen on congenital liver Insr knockout*(21)*. Wild-type (WT) or mutant human INSR, or GFP alone, were then expressed in knockout liver by injection *via* the tail vein at 10 weeks of age of adenovirus (AdV) containing transgenes under the control of the liver specific albumin promoter, to create a series of “add back” models of human receptoropathy, denoted here by L-IRKO+WT, L- IRKO+(*INSR mutation*) and L-IRKO+GFP respectively. All *INSR* constructs included C-terminal myc-tags to aid detection of transgene expression. Liver wild-type (L-WT) mice with unperturbed liver *Insr* expression were generated by injecting *Insr*^*loxP/loxP*^ mice with AAV encoding GFP at 8-weeks of age, followed by AdV encoding GFP at 10-weeks of age, and served as additional controls (Figure 1A).

**Figure 1.**
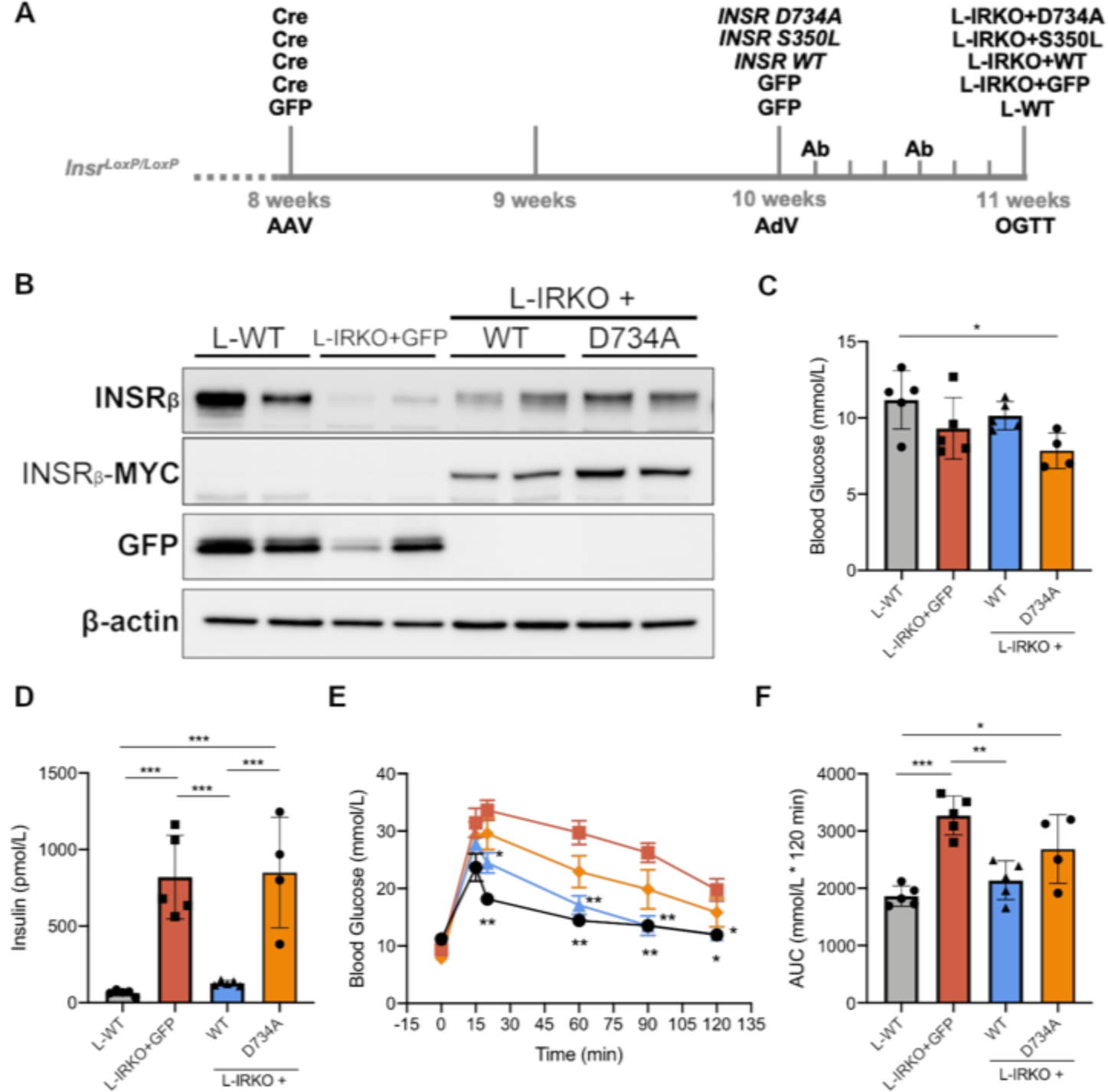
Hyperglycemia and insulin resistance due to liver insulin receptor deletion are rescued by WT but not mutant human INSR. (**A**) Schematic representation of the generation of the insulin receptoropathy model. (**B**) Western blot of liver lysates from mice on completion of oral glucose tolerance test (OGTT) and probed for insulin receptor β subunit (INSR_β_), MYC, GFP or β-actin. (**C**) Blood glucose and (**D**) insulin concentrations in mice after 5 h fasting. (**E**) Results of OGTT (2g/kg glucose) after 5 h fasting, L-IRKO+GFP (squares), L-IRKO+D734A (diamonds), L-IRKO+WT (upward triangles), L-WT (circles). (**F**) OGTT areas under the curve. L-WT mice = AAV-GFP/AdV-GFP (i.e. GFP control only), L-IRKO+GFP mice = AAV-iCre/AdV-GFP (i.e. liver *Insr* knockout only), L-IRKO+WT = AAV-iCre/AdV-Hs*INSR*-WT-myc (i.e. L-IRKO with WT INSR add back), L-IRKO+D734A = AAV-iCre/AdV-Hs*INSR*-D734A-myc (i.e. L-IRKO with D734A INSR add back). Data in **C, D** and **F** are shown as mean ± SD, with statistical significance tested by one-way ANOVA with Tukey’s multiple comparison test. n = 5 per group, except L-IRKO+D734A (n = 4). * p<0.05, ** p<0.01, *** p<0.001. Data in **E** are mean ± SEM, with statistical significance of difference from L-IRKO+GFP tested by two-way repeated measures ANOVA with Tukey’s multiple comparisons test. * p<0.05, ** p<0.01.

Two INSR mutations, D734A and S350L, were selected for study based on prior evaluation in cell signaling assays*(18)*. Both D734A*(22)* and S350L*(23)* mutations produce receptors that are normally processed and expressed at the cell surface but demonstrate severely reduced insulin binding and autophosphorylation. Indeed the INSR D734A mutation lies in the αCT segment of the extracellular domain of the INSR, which is a critical structural components of insulin binding site 1, initially identified in biochemical studies*(24)*. Importantly, both mutant INSR have been shown to be activatable by anti-INSR antibodies*(18, 25)*.

The INSR D734A mutant was used first to evaluate the “add back” approach. Western blots of liver lysates confirmed efficient deletion of endogenous *Insr* in mice administered AAV-Cre, expression of GFP in mice administered control virus, and expression of myc-tagged INSR in mice administered AdV encoding human *INSR* transgenes (Figure 1B). Blood glucose concentrations following 5 h fasting were the same across all groups except L-IRKO+D734A, which demonstrated decreased fasting blood glucose (Figure 1C), presumably due to the action of increased insulin concentration on unaffected peripheral tissues (Figure 1D). L-IRKO+GFP mice demonstrated marked increase of fasting blood insulin concentration compared to L-WT mice (p<0.001), and this was rescued upon expression of human WT INSR in the liver (p<0.001) (L-IRKO+WT) (Figure 1D). Likewise, mice expressing the INSR D734A mutant demonstrated significant elevation of blood insulin concentration compared to L-WT and L-IRKO+WT (both p<0.001) (Figure 1D). L-IRKO+GFP mice were more glucose intolerant than L-WT mice (Figure 1E) with increased glucose excursion during 120min OGTT, assessed as area under the curve (Figure 1F). Add-back of human WT INSR but not mutant D734A INSR restored glucose tolerance (Figure 1F). These findings confirmed that an “add back” model of human insulin receptoropathy was capable of discriminating clearly between WT and mutant INSR.

### Antibody treatment down-regulates wild-type human INSR expression with minimal effect on glucose homeostasis

The primary aim of this study was to assess whether dysfunctional, mutant INSR can be activated by anti-INSR antibodies. However, such bivalent antibodies can also bind and activate WT human INSR*(14)*, and naturally occurring polyclonal anti-INSR antibodies induce hypoglycaemia at low titres, and extreme insulin resistance at high titres in humans*(19)*. On the other hand, monoclonal human anti-INSR antibodies have been suggested as a therapeutic strategy in common forms of diabetes, without evaluation of effects on receptor expression*(9–12)*. Understanding the balance between surrogate agonism and receptor downregulation is likely to be a critical consideration in development of antibody therapeutics for receptoropathy. The effects of anti-INSR antibodies were thus first assessed in mice with WT INSR added back. Antibody was administered at a dose of 10mg/kg, delivered by intraperitoneal injection at 1 and 4 days after adenoviral injection, before metabolic evaluation 7 days after adenoviral injection (Figure 1A). Antibody dose and treatment schedule were based on previous studies of agonistic INSR antibodies*(9–11)*. Control antibody-treated animals demonstrated a similar pattern of glucose tolerance and circulating insulin concentrations to those in initial characterization studies (Supplementary Figure 1). Anti-INSR antibodies caused significant (p<0.01) down-regulation of myc-tagged INSR protein expression (Figure 2A and B) with no change in mRNA expression of human *INSR* transgene (Figure 2C) amongst treatment groups, indicating that the decrease in myc-tagged INSR protein levels was not due to failed liver transduction with transgene. No further decrease in the very low level of residual endogenous *Insr* was seen (Figure 2D). Treatment of L-IRKO+WT mice with anti-INSR antibodies did not alter glucose tolerance (Figure 2E and F), but fasting blood glucose concentration was mildly decreased in 83-14-treated mice compared to control and 83-7 treated mice (p<0.01 and p<0.05 respectively) (Figure 2G). Fasting blood insulin concentration was unaffected by either antibody treatment (Figure 2H).

**Figure 2.**
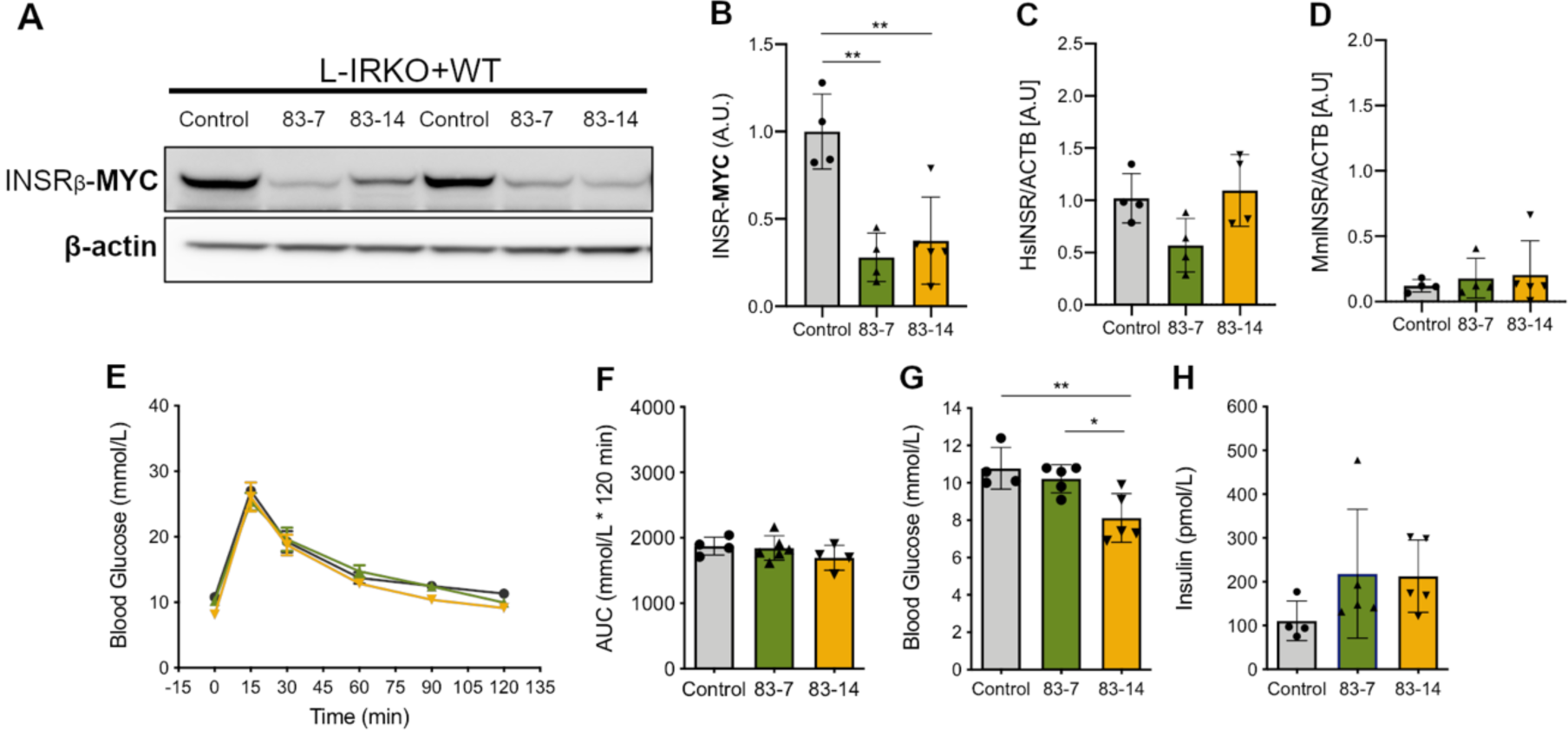
Antibody treatment down-regulates wild-type human INSR expression in mouse liver with minimal effect on glucose homeostasis. L-IRKO + WT mice were dosed twice over one week with 10mg/kg control (n=4) or anti-INSR antibodies 83-7 (n=5) or 83-14 (n=5) as indicated. (**A**) Western blot of liver lysates from L-IRKO+WT mice at the completion of OGTT, probing for MYC-tagged β subunit or β-actin as indicated. Quantification of (**B**) Myc-tagged human INSR protein, (**C**) human *INSR* mRNA, and (**D**) endogenous *Insr* mRNA in livers from the same experiment. mRNA was quantified by qPCR. (**E**) OGTT (2g glucose/kg) after 5 h fast in antibody treated L-IRKO+WT mice. (**F**) Area under blood glucose curves during OGTT in antibody treated L-IRKO+WT mice. (**G**) Blood glucose concentrations in antibody treated L-IRKO+WT mice after 5 h fast. (**H**) Insulin concentrations in antibody treated L-IRKO+WT mice after 5 h fast. All data (except **E**) are shown as mean ± SD, with statistical significance tested by one-way ANOVA with Tukey’s multiple comparison test, * p<0.05, ** p<0.01. In (**E**) data shown are mean ± SEM. Circles = control antibody. Upward triangles = 83-7 antibody. Downward triangles = 83-14 antibody. Lack of statistical significance was determined by two-way repeated measures ANOVA with Tukey’s multiple comparisons test.

As the anti-INSR antibodies used do not bind mouse Insr, off target metabolic effects were not anticipated. To confirm this anti-INSR antibodies were administered to L-WT mice (Supplementary Figure 2). As expected, no effect on endogenous liver Insr protein expression (Supplementary Figure 2A and B), mRNA expression (Supplementary Figure 2C), glucose tolerance (Supplementary Figure 2D and E), fasting blood glucose concentration (Supplementary Figure 2F) or fasting blood insulin concentration (Supplementary Figure 2G) was seen. L-IRKO+GFP mice, with severely reduced liver Insr expression (Supplementary Figure 2A, H and I), also showed no change in any metabolic assessment (Supplementary Figure 2J-M).

### Antibody treatment improves glucose tolerance and hyperinsulinemia in receptoropathy models, but down-regulates INSR protein expression

In L-IRKO+GFP mice with ‘add-back’ of INSR D734A, treatment with 83-7 and 83-14 antibodies downregulated myc-tagged INSR protein levels compared to control-treated animals (p<0.0001) (Figure 3 A and B). This was not accompanied by any change in either human *INSR* transgene mRNA (Figure 3C) or endogenous mouse *Insr* mRNA (Figure 3D). Despite this, treatment with 83-7 and 83-14 did improve glucose tolerance (Figure 3E and F). This was not accompanied by any change in fasting blood glucose concentrations (Figure 3G), however antibody 83-14 significantly (p<0.05) reduced fasting insulin concentrations in L-IRKO+D734A animals (Figure 3H).

**Figure 3.**
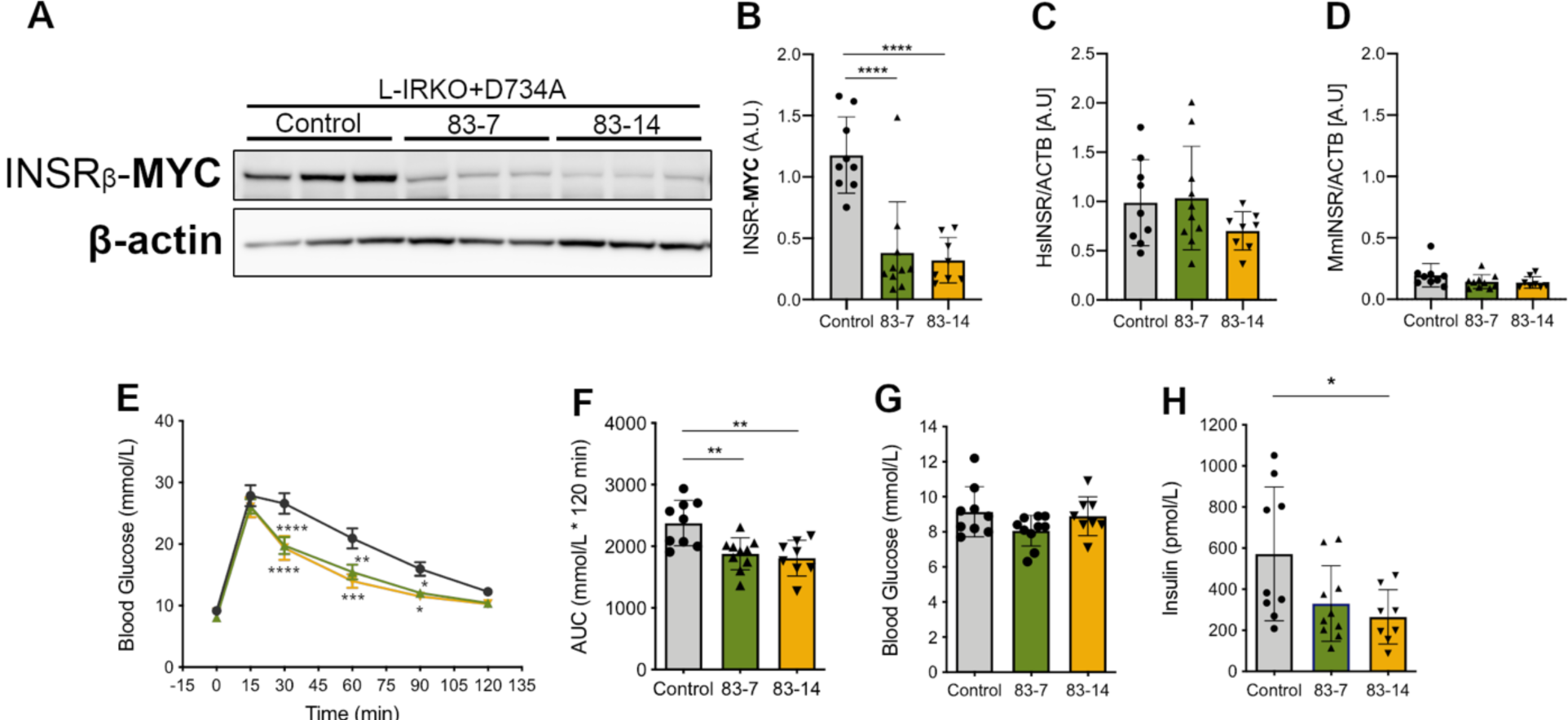
Antibody treatment improves glucose tolerance and hyperinsulinemia in INSR D734A add-back mice, but down-regulates INSR protein expression. L-IRKO+D734A mice were treated twice over one week with 10mg/kg control (n=9) or anti-INSR 83-7 (n=10) or 83-14 (n=8) antibodies. (**A**) Western blot of liver lysates from L-IRKO+D734A mice at completion of OGTT, probed for the proteins as indicated. Quantification of (**B**) Myc-tagged human INSR protein, (**C**) human *INSR* mRNA, and (**D**) endogenous *Insr* mRNA in livers from the same experiment. mRNA was quantified by qPCR. (**E**) OGTT (2g glucose/kg) after 5 h fast in antibody treated L-IRKO+D734Amice. (**F**) Area under blood glucose curves during OGTT in antibody treated L-IRKO+D734A mice. (**G**) Blood glucose concentrations in antibody treated L-IRKO+D734A mice after 5 h fast. (**H**) Insulin concentrations in antibody treated L-IRKO+D734A mice after 5 h fast. All data (except **E**) are shown as mean ± SD, with statistical significance tested by one-way ANOVA with Tukey’s multiple comparison test. In (**E**) data shown are mean ± SEM. Circles = control antibody. Upward triangles = 83-7 antibody. Downward triangles = 83- 14 antibody. Statistical significance was tested by two-way repeated measures ANOVA with Tukey’s multiple comparisons test. * p<0.05, ** p<0.01, *** p<0.001, ****p<0.0001.

In L-IRKO+GFP mice with ‘add-back’ of S350L mutant human INSR, treatment with 83-7 and 83-14 also reduced myc-tagged INSR protein levels (Figure 4 A and B). As in L-IRKO+WT and L-IRKO+D734A mice, this was not due to failed liver transduction with human *INSR*, as qPCR demonstrated stable human *INSR* transgene mRNA across all treatment conditions (Figure 4C) and effective deletion of endogenous mouse *Insr* (Figure 4D). Animals treated with 83-7 and 83-14 showed only a trend to improved glucose tolerance (Figure 4 E and F), and neither antibody lowered fasting blood glucose concentrations (Figure 4G). Treatment of L-IRKO+S350L mice with anti-INSR antibody 83-7 did reduce fasting blood insulin concentration compared to control and 83-14-treated animals (both p<0.05), indirectly demonstrating hypoglycaemic action of antibody (Figure 4H). Collectively these findings demonstrate that anti-INSR monoclonal antibodies improve glucose tolerance and reduce fasting hyperinsulinemia in mice expressing human INSR mutations that cause recessive insulin receptoropathy, but that the magnitude of the improvement seen is likely attenuated by downregulation of INSR expression.

**Figure 4.**
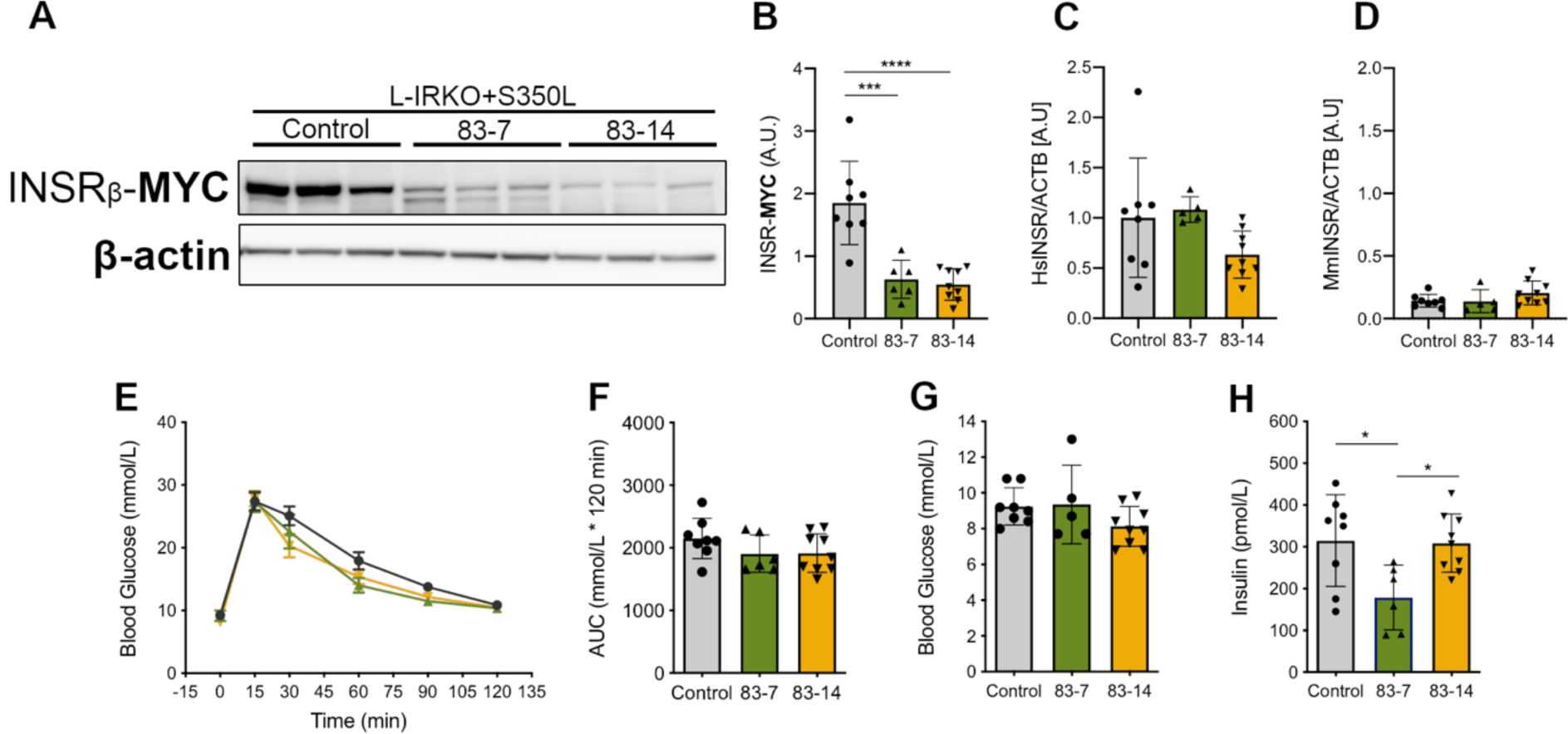
Antibody treatment reduces fasting hyperinsulinemia in INSR S350L add-back mice but downregulates INSR protein expression. L-IRKO+S350L mice were treated twice over one week with 10mg/kg control (n=8) or anti-INSR 83-7 (n=6) or 83-14 (n=9) antibodies. (**A**) Western blot of liver lysates from mice at completion of OGTT were probed for the proteins indicated. Quantification of (**B**) Myc-tagged human INSR protein, (**C**) human *INSR* mRNA and (**D**) endogenous *Insr* mRNA in livers from the same experiment. mRNA was quantified by qPCR. (**E**) OGTT (2g glucose/kg) after 5 h fast in antibody treated L-IRKO+S350L mice. (**F**) Area under blood glucose curves during OGTT in antibody treated L-IRKO+S350L mice. (**G**) Blood glucose concentrations in antibody treated L-IRKO+S350L mice after 5 h fast. (**H**) Insulin concentrations in antibody treated L-IRKO+S350Lmice after 5 h fast. All data (except **E**) are shown as mean ± SD, with statistical significance tested by one-way ANOVA with Tukey’s multiple comparison test. In (**E**) data shown are mean ± SEM. Circles = control antibody. Upward triangles = 83-7 antibody. Downward triangles = 83-14 antibody. Lack of statistical significance was determined by two-way repeated measures ANOVA with Tukey’s multiple comparisons test. * p<0.05, ** p<0.01, *** p<0.001, ****p<0.0001.

## Discussion

Extreme congenital IR was first clinically described as Donohue Syndrome, and the less severe Rabson Mendenhall Syndrome (RMS), decades before the insulin receptor was identified, and thus long before the genetic cause, namely bi-allelic *INSR* mutations, was established*(26, 27)*. Both syndromes feature extreme metabolic derangement, characterised by high blood glucose concentration that is unresponsive or minimally responsive to insulin therapy. They also feature severely impaired linear growth and underdevelopment of insulin-responsive tissues such as skeletal muscle and adipose tissue. Less intuitively, marked overgrowth of other tissues and organs including skin, kidneys, liver, gonads and colonic mucosa is also seen, and may pose clinical challenges*(2)*. Overgrowth is thought to be driven by compensatory elevation of blood insulin concentration, which can act on the trophic insulin-like growth factor 1 (IGF1) receptor, which is structurally similar to the INSR*(2)*.

The clinical course of recessive insulin receptoropathy is bleak, with death common between infancy, at which stage it often occurs during viral infection, and early adolescence, when it is more likely due to complications of uncontrolled diabetes such as ketoacidosis or microvascular damage. There is a major unmet therapeutic need for novel insulin-mimetic agents, ideally with no action on the IGF1 receptor. Genetic considerations suggest that only a small degree of activation of ‘non-functional’ receptors may be required to achieve major clinical benefits: Donohue syndrome is caused by complete or near complete loss of receptor function, while Rabson Mendenhall syndrome, with a better prognosis, features around 10-20% receptor function. Autosomal dominant insulin receptoropathy, which usually presents only around puberty, features no more than 25% receptor function, while lack of one INSR allele (50% function) has not been associated with IR. This suggests a steep relationship between INSR function and prognosis between 0 and 25% receptor function.

We previously demonstrated the ability of bivalent, specific anti-INSR antibodies to act as surrogate ligands on a series of mutant INSR in cell culture models*(18)*, and now report their evaluation *in vivo* in a novel mouse model of human insulin receptoropathy. The ‘humanised’ mouse model of insulin receptoropathy was generated by using sequential viral infection to knock out endogenous *Insr* and then to re-express human *INSR*. This enabled changes in metabolic outcomes upon antibody treatment to be attributed to action on re-expressed human mutant INSR as the monoclonal anti-INSR antibodies tested do not bind rodent Insr*(14)*. Use of a viral strategy made liver the most tractable organ to target, and also had the benefit that liver parenchyma is particularly accessible to antibody due to the fenestration of hepatic capillaries. This approach also avoided the compensatory responses reported in congenital liver Insr deficiency*(21)*, while offering flexibility to study various mutant human INSR transgenes without generating distinct genetically modified mouse lines. On the other hand technical success relies on efficient administration of viral vectors by skilled operators, and AdV vectors limit duration of transgene expression, constraining the time window for study.

Encouragingly, both monoclonal anti-INSR Abs tested - 83-7 and 83-14 - did improve glucose tolerance in L-IRKO+D734A mice, while 83-14 treatment also lowered fasting blood insulin concentration in these mice (Figure 3H). 83-7 lowered fasting blood insulin concentration in L-IRKO+S350L mice (Figure 4H). Collectively, these observations demonstrate that anti-INSR antibodies can improve glucose tolerance and reduce fasting hyperinsulinemia in mice expressing human INSR that cause severe disease in humans, adding to evidence that antibody-based surrogate agonism may be of metabolic benefit *in vivo*. The effects observed in this acute receptoropathy model were modest and not fully consistent between mutants or antibodies, or indices of IR. However, several factors may have adversely affected the potential for antibodies to ameliorate the condition. First, overexpression of the human INSR added back may have attenuated the degree of IR that mutants confer compared to humans with endogenous expression of the same mutations, reducing the dynamic range of IR of the model. This may explain the relatively mild IR seen with S350L at baseline (Figure 4), despite this mutation being found to cause RMS in several unrelated families. While future calibration of the viral models described against mice with endogenous expression of mutant receptors would be of great interest, the need to study human INSR rather than mouse Insr makes this a challenging technical proposition.

A second potential reason why metabolic effects of antibodies were not larger has more profound implications for INSR surrogate agonist-based strategies for treating IR. Antibody treatment downregulated receptor expression across all INSR species studied, as expected from the known coupling of receptor activation to internalisation. Following internalisation by endocytosis receptors are trafficked through the endosomal/lysosomal pathway and either recycled to the cell surface in the unliganded state or degraded*(28)*. The mechanisms governing internalisation, trafficking, and the balance of subsequent recycling and degradation in response to stimulation are poorly understood, but the potential importance of this in the context of anti-INSR antibodies is known from studies of Type B insulin resistance*(19)*. This is a naturally occurring, acquired form of insulin receptoropathy driven by anti-INSR antibodies. It is well known that low titres of such antibodies can produce clinically important hypoglycaemia, but that when antibody titres rise severe receptor downregulation and fulminant IR occurs that may be life threatening*(19)*. This harmful effect of high antibody levels will likely narrow the therapeutic window for agonistic anti-INSR antibodies in recessive receptoropathy unless ways of modulating internalisation and/or receptor recycling and degradation rates are devised.

In summary, we report a novel approach to modelling recessive human insulin receptor defects in the mouse using sequential virally-mediated knockout of endogenous and re-expression of human insulin receptor. This yielded mice with acute IR due to two previously studied INSR mutations that have been shown in cell models to exhibit activation by anti-INSR antibodies. Injection of well characterised monoclonal anti-INSR antibodies improved IR in both models, however the magnitude of the effect is likely to have been limited by downregulation of receptor. Our findings confirm the potential utility of surrogate agonist strategies for treating lethal insulin receptoropathy, but caution that receptor downregulation may attenuate the benefits realised unless this can concomitantly be reduced.

## Materials and Methods

### Mice

All mouse experiments were approved under the UK Home Office Animals (Scientific Procedures) Act 1986 following ethical review by the University of Cambridge Animal Welfare and Ethical Review Board. *Insr*^*loxP/loxP*^ mice were described previously*(21)*, as was use of adeno-associated virus to deliver *Cre* to generate liver insulin receptor knock out (L-IRKO) mice*(20). Insr*^*loxP/loxP*^ mice were purchased from The Jackson Laboratories (Bar Harbor, ME, USA). Mice were fed regular laboratory SAFE105 chow diet (Safe Diets, Augy, France) throughout the study. Male mice were injected *via* the tail-vein at 8 weeks of age with 10^11^ copies per mouse of adeno-associated virus serotype 8 containing a hybrid promoter based on the thyroid hormone-binding globulin (TBG) promoter and macroglobulin/bikunin enhancer. This permitted liver-specific expression of iCre or eGFP to generate L-IRKO or liver wild-type (L-WT) mice, respectively. At 10 weeks of age, mice were injected *via* the tail-vein with 5×10^9^ infectious units (IFU) per mouse of adenovirus serotype 5 containing the liver albumin promoter driving liver-specific expression of either c-terminal myc-tagged human insulin receptor WT, S350L, D734A mutants, or eGFP (L-IRKO+GFP and L-WT). Mice were treated twice over one week with 10mg/kg antibody via intraperitoneal injection, with the first dose given the day after AdV was administered. Experiments were performed a week after adenovirus injection. Mice were euthanised by cervical dislocation at the conclusion of the oral glucose tolerance test and tissues were harvested and snap-frozen immediately in liquid nitrogen before storage at −80°C until further processing.

### Metabolic measurements

For oral glucose tolerance tests (OGTT), mice were fasted for 5 h prior to oral gavage of glucose at 2g/kg body weight. Blood glucose measurements were made using a blood glucose analyser (AlphaTRAK) at 0, 15, 30, 60, 90, 120 minutes. For plasma insulin analysis, tail blood was collected at 0 and 15 min into glass micro-haematocrit capillary tubes with sodium heparin (Hirshmann-Laborgeräte, Germany). Insulin concentrations were determined by a sandwich immunoassay providing an electrochemiluminescent readout (Meso Scale Discovery, Maryland, USA).

### Liver protein extraction and Western blotting

Liver tissues were homogenised in RIPA buffer (Thermo Fisher Scientific, MA, USA) containing protease and phosphatase inhibitors (Roche, Penzberg, Germany) using Matrix D ceramic beads and a FastPrep-24™ benchtop homogeniser (MP Biomedicals, CA, USA). Lysates were cleared of insoluble debris by centrifugation prior to determination of protein concentration by BCA assay (BioRad, CA, USA). Lysates were electrophoresed through either 4-12% NuPAGE or 8% E-PAGE gels (Thermo Fisher Scientific, MA, USA) and transferred to nitrocellulose by iBlot 2 dry blotting system (Thermo Fisher Scientific, MA, USA). The following antibodies were used for immunoblotting at a dilution of 1:1000: INSR (3025), Myc-tag (2276), Beta-actin (4967) from Cell Signaling Technology (MA, USA), and at a dilution of 1:2000: GFP (ab290) from Abcam (Cambridge, UK). Horseradish peroxidase (HRP)-conjugated secondary antibodies and Immobilon Western Chemiluminescent HRP substrate (Millipore, Massachusetts, USA) were used to detect protein-antibody complexes, and grey-scale 16-bit TIFs captured with an ImageQuant LAS4000 camera system (GE Healthcare Lifesciences, Massachusetts, USA). Pixel density of grey-scale 16-bit TIFs was determined in ImageJ 1.52b (NIH, USA). The rectangle tool was used to select lanes and the line tool to enclose the peak of interest and subtract background. The magic-wand tool was used to select the peak area and obtain the raw densitometry value. Normalised values for INSR were scaled to the mean expression of INSR in L-WT tissues and Myc-tagged INSR values were scaled to the mean expression of INSR-myc in control antibody treated L-IRKO+WT animals.

### Liver mRNA isolation and quantitative PCR

Total RNA was isolated from liver tissues using the Direct-zol RNA extraction kit from Zymo Research (Irvine, CA, USA). Complementary DNA was reverse-transcribed using Moloney murine leukemia virus reverse transcriptase (Promega, Wisconsin, USA). Relative expression of genes of interest were quantified by real-time PCR using TaqMan™ gene expression assays (Mm02619580_g1 ACTB, Mm01211877_m1 Mm Insr, Hs00961560_m1 Hs INSR) and the QuantStudio™ 7 Flex Real-Time PCR system (Thermo Fisher Scientific, MA, USA) and results analysed by the comparative Ct method. Validation experiments were performed both to confirm species specificity of TaqMan™ gene expression assays and that their relative amplification efficiencies permitted analysis by the comparative Ct method.

### Statistical analysis

Mice were randomly assigned to viral injection schedules and antibody treatment groups. Investigators were blind to the assignment at the time of administering treatments and performing experiments. All data presented are the mean ± S.D with the exception of the OGTT histograms which are the mean ± S.E.M. Statistical analysis were performed using GraphPad Prism 8 for macOS version 8.3.0 (GraphPad Software, CA, USA). Protein abundance, mRNA expression, fasting blood glucose and insulin, and AUC (blood glucose mmol/L*120min) were computed using the trapezoid rule and analysed by one-way ANOVA followed by Tukey’s multiple comparisons test. Blood glucose levels during OGTT were analysed by two-way (repeated measures) ANOVA followed by Tukey’s multiple comparisons test. A probability level of 5% (p<0.05) was considered statistically significant.

## Supplementary Materials

**Supplementary Figure 1.**
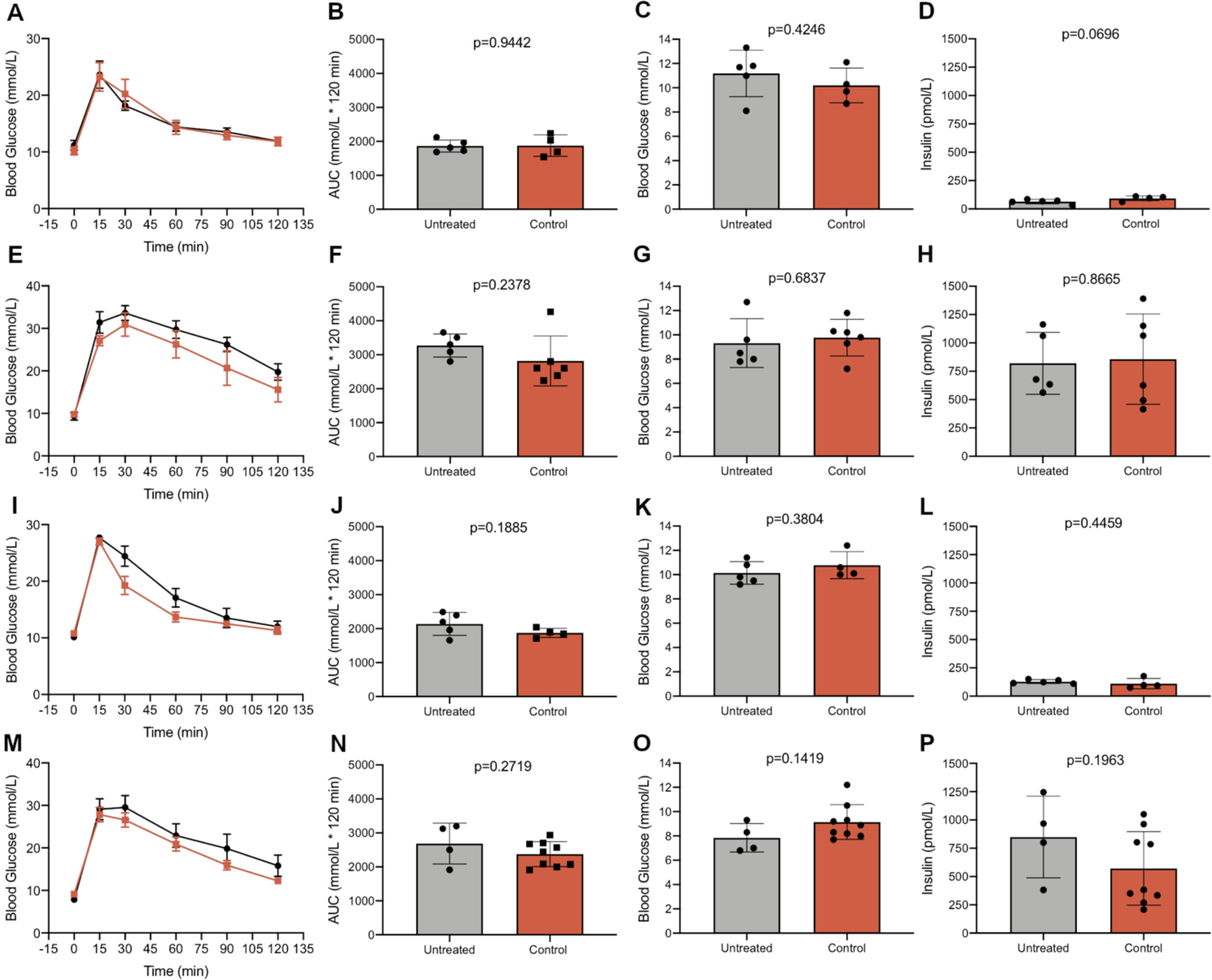
Control antibody has no metabolic effect in mice with acute liver insulin receptoropathy. (**A - D**) L-WT mice, (**E – H**) L-IRKO+GFP mice, (**I – L**) L-IRKO+WT mice, (**M – P**) L- IRKO+D734A mice. Results of OGTT (2g/kg glucose) after 5 h fasting (**A, E, I, M**) and OGTT areas under the curve (**B, F, J, N**). (**C, G, K, O**) Blood glucose and (**D, H, L, P**) insulin concentrations in mice after 5 h fasting. L-WT mice = AAV-GFP/AdV-GFP (i.e. GFP control only), L-IRKO+GFP mice = AAV-iCre/AdV-GFP (i.e. liver *Insr* knockout only), L-IRKO + WT = AAV-iCre/AdV-Hs*INSR*-WT-myc (i.e. L-IRKO with WT INSR add back), L- IRKO + D734A = AAV-iCre/AdV-Hs*INSR*-D734A-myc (i.e. L-IRKO with D734A INSR add back). Data in **A, E, I, M** are shown are means ± SEM, with statistical significance of difference from L-IRKO+GFP tested by two-way repeated measures ANOVA with Sidak’s multiple comparisons test. All other data are shown as mean ± SD, with statistical significance determined by unpaired two-tailed t-test.

**Supplementary Figure 2.**
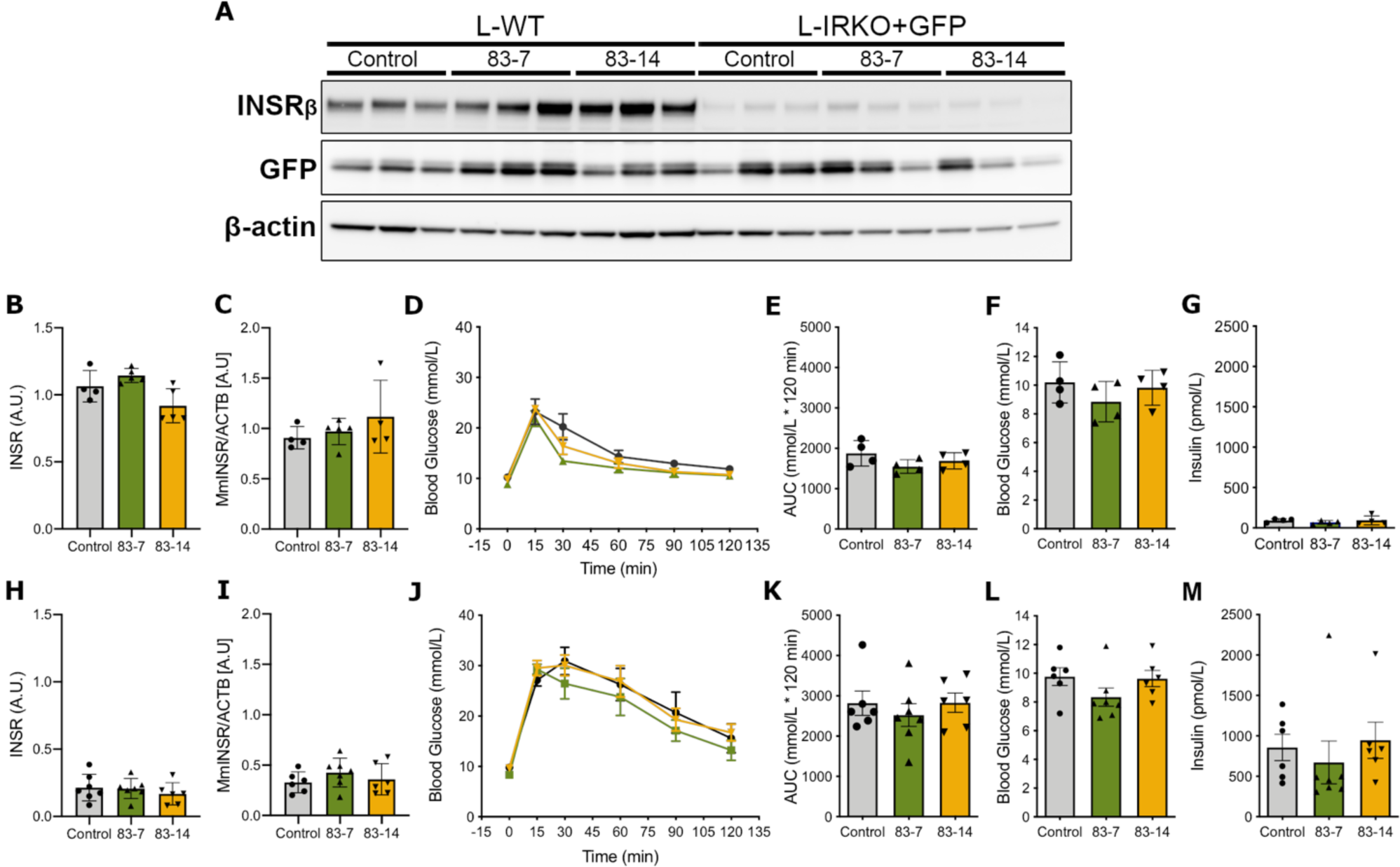
Antibody treatment has no effect in mice not expressing human INSR. L-WT (**B** - **G**) and L-IRKO+GFP (**H**- **M**) mice were treated twice over the course of a week with 10mg/kg control or anti-INSR antibodies 83-7 or 83-14 as indicated. (**A**) Representative Western blot of lysates from livers harvested from L-WT and L-IRKO+GFP mice at the completion of oral glucose tolerance test (OGTT) and probed for specific proteins as indicated. INSR protein expression (**B, H**) and *Insr* gene expression (**C, I**) in antibody treated L-WT and L-IRKO+GFP mice, respectively. Glucose tolerance test 2g/kg administered by oral gavage after 5 h fast, in antibody treated L-WT (**D**) and L- IRKO+GFP (**J**). Data are mean ± SEM. Circles = control antibody. Upward triangles = 83-7 antibody. Downward triangles = 83-14 antibody. Lack of statistical significance determined by two-way repeated measures ANOVA with Tukey’s multiple comparisons test. Cumulative measurement of blood glucose during 120 min OGTT in antibody treated L-WT (**E**) and L-IRKO+GFP (**K**) mice. Blood glucose concentrations in antibody treated L-WT (**F**) and L-IRKO+GFP (**L**) mice after 5 h fast. Insulin concentrations in antibody treated L-WT (**G**) and L-IRKO+GFP (**M**) mice after 5 h fast. All data (except D and J) are mean ± SD, lack of statistical significance was determined by one-way ANOVA with Tukey’s multiple comparison test. L-WT n = 4 per group. L-IRKO+GFP n = 6 per group control and 83-7 treated animals and n = 7 for the 83-14 treated group.

## Acknowledgments

We are grateful to Allie Finigan (Department of Medicine, University of Cambridge) for assistance in performing IV injections, to Amy Warner and Daniel Hart (Disease Model Core, MRC Metabolic Diseases Unit (MDU), University of Cambridge), Keith Burling (Clinical Biochemistry Assay Laboratory, University of Cambridge) and James Warner (Histology Core, MRC MDU) for technical assistance.

## Author contributions

Conceptualization: GVB, KS, RKS; Formal Analysis: GVB, KS, RKS; Performed investigations: GVB, HW, ER, SG; Writing: GVB, KS, RKS; Project Administration: GVB, KS, RKS; Supervision: GVB, KS, RKS.

## Data and materials availability

The datasets generated during and/or analysed during the current study are available from the corresponding author on reasonable request

## Tables

This manuscript does not contain any tables

